# BraCeR: Reconstruction of B-cell receptor sequences and clonality inference from single-cell RNA-sequencing

**DOI:** 10.1101/185504

**Authors:** Ida Lindeman, Guy Emerton, Ludvig M. Sollid, Sarah A. Teichmann, Michael J.T. Stubbington

**Affiliations:** Centre for Immune Regulation and Department of Immunology, University of Oslo and Oslo University Hospital, 0372 Oslo, Norway.; Wellcome Trust Sanger Institute, Hinxton, Cambridgeshire, CB10 1RQ, United Kingdom.; KG Jebsen Coeliac Disease Research Centre, University of Oslo, 0372 Oslo, Norway.; Theory of Condensed Matter, Cavendish Laboratory, 19 JJ Thomson Ave, Cambridge CB3 0HE, United Kingdom.

## Abstract

Reconstruction of antigen receptor sequences from single-cell RNA-sequencing (scRNA-seq) data allows the linking of antigen receptor usage to the full transcriptomic identity of individual B lymphocytes, without having to perform additional targeted repertoire sequencing (Rep-seq). Here we report BraCeR (freely available at https://github.com/teichlab/bracer/), an extension of TraCeR [1], for reconstruction of paired full-length B-cell receptor sequences and inference of clonality from scRNA-seq data (**Supplementary Note 1**).

---

BraCeR builds on the well-verified pipeline of TraCeR for assembly of BCR sequences from paired-end or single-end reads, modified to account for somatic hypermutations (SHM) and isotype switching (**Supplementary Notes 2** and **3**). In addition, a ‘Build’ mode facilitates the creation of resource files for the analysis of species beyond human and mouse.

BraCeR was tested against experimental human and mouse scRNA-seq data with various SHM rates and repertoire diversities [2, 3]. The reconstruction accuracy was similar to that of BASIC, a previously reported tool for BCR reconstruction [2]. Comparatively, BraCeR yielded a reconstruction efficiency that was somewhat superior for the long (125 bp) reads and similar to
BASIC for short (50 bp) reads (**Supplementary Note 4** and **Supplementary Tables 2-4**). Importantly however, BraCeR allows for reconstruction of additional heavy and light chains present in a cell, and it can identify non-productively rearranged chains. It is also possible to provide BraCeR with FASTA files containing BCR sequences assembled by other means or determined by single-cell Rep-seq for downstream analyses.

BraCeR further improves upon BASIC by using the reconstructed sequences to infer clonal relationships and perform immunoglobulin lineage reconstruction. BraCeR identifies the isotype of each reconstructed BCR and collapses highly similar reconstructed sequences in a cell. It then quantifies the expression of the BCR sequences and identifies the most highly expressed chains for each locus in each cell. Potential cell multiplets or cross-cell contaminations are reported, and are by default removed from downstream analyses (**Supplementary Note 5**).

Clonally related cells are grouped into clonotypes based on the reconstructed chains in each cell. Productively rearranged reconstructed BCRs for each locus are grouped into clones based on V- and J-gene assignments using custom Python scripts and a previously described method for comparison of CDR3 sequence similarity [4] (**Supplementary Note 6**). Clonal networks are constructed based on shared clonal heavy and light chains and then visualised (**Fig. 1A**). As an optional feature, BraCeR can also use existing tools [4-6] to construct lineage trees for each group of clonal cells (**Fig. 1B**), with the inferred germline sequence as outgroup, facilitating linkage of transcriptomic phenotype to the evolution of immunoglobulin sequences (**Supplementary Note 7**).

**Figure 1.**
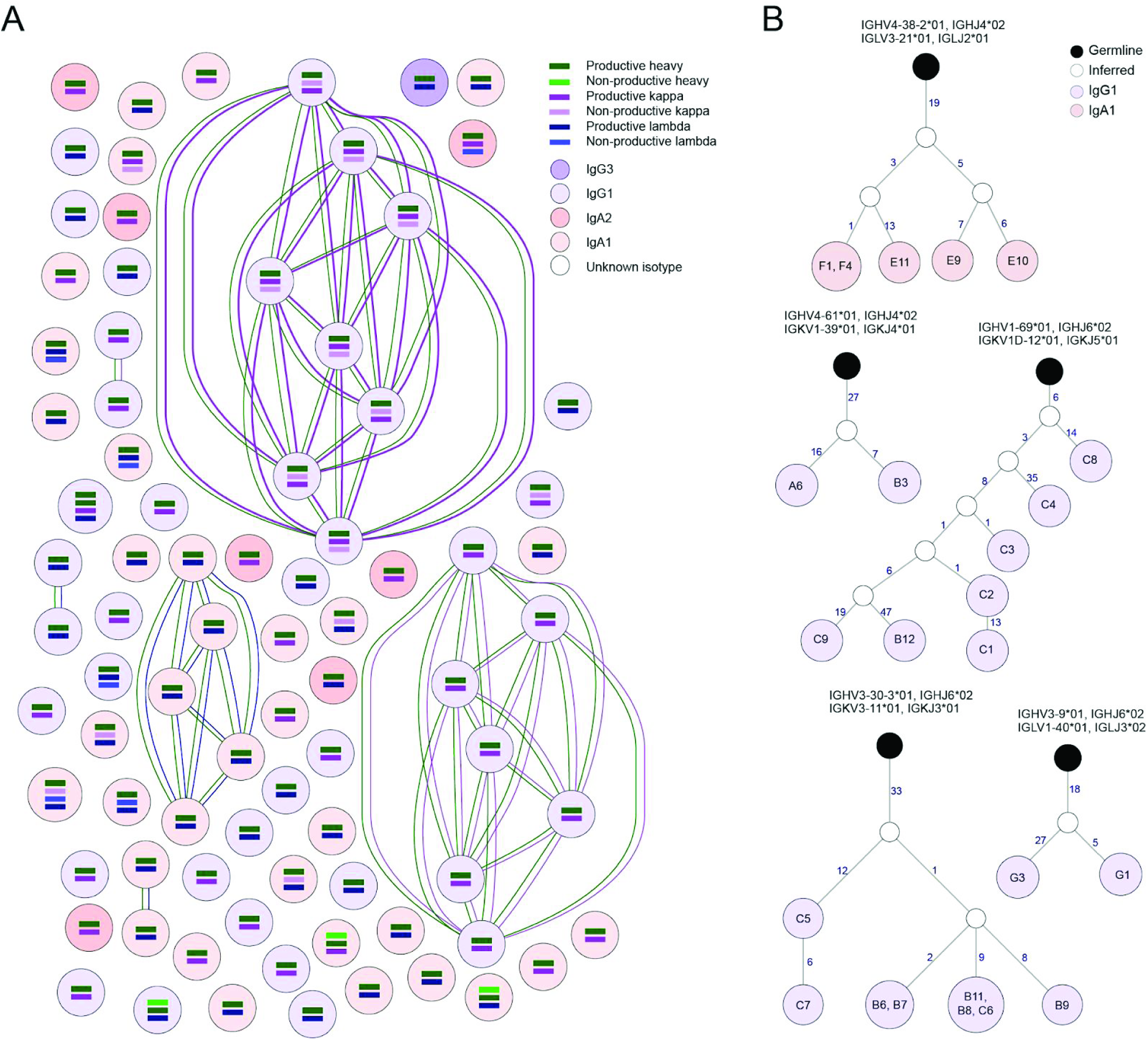
Clonal network (**A**) and lineage trees (**B**) for each clonotype defined by BraCeR for a human plasmablast scRNA-seq dataset [2].

With an easy-to-use command-line interface, BraCeR provides a complete pipeline for clonal inference and lineage tracing of B cells; raw scRNA-seq reads can be processed all the way to clonal networks and lineage trees. Additional output includes summary files and graphs summarising reconstructed sequences and isotype usage. BraCeR also creates tab-delimited database files compatible with the Immcantation portal tool suite following the standards of the Adaptive Immune Receptor Repertoire (AIRR) Community, thus facilitating further analyses of the reconstructed BCR sequences.

## Acknowledgements

We would like to thank Omri Snir, Shuo-Wang Qiao and Steven Kleinstein for helpful discussions.

## Supplementary information

### Supplementary Note 1. BraCeR pipeline overview

**Supplementary Note 1.**
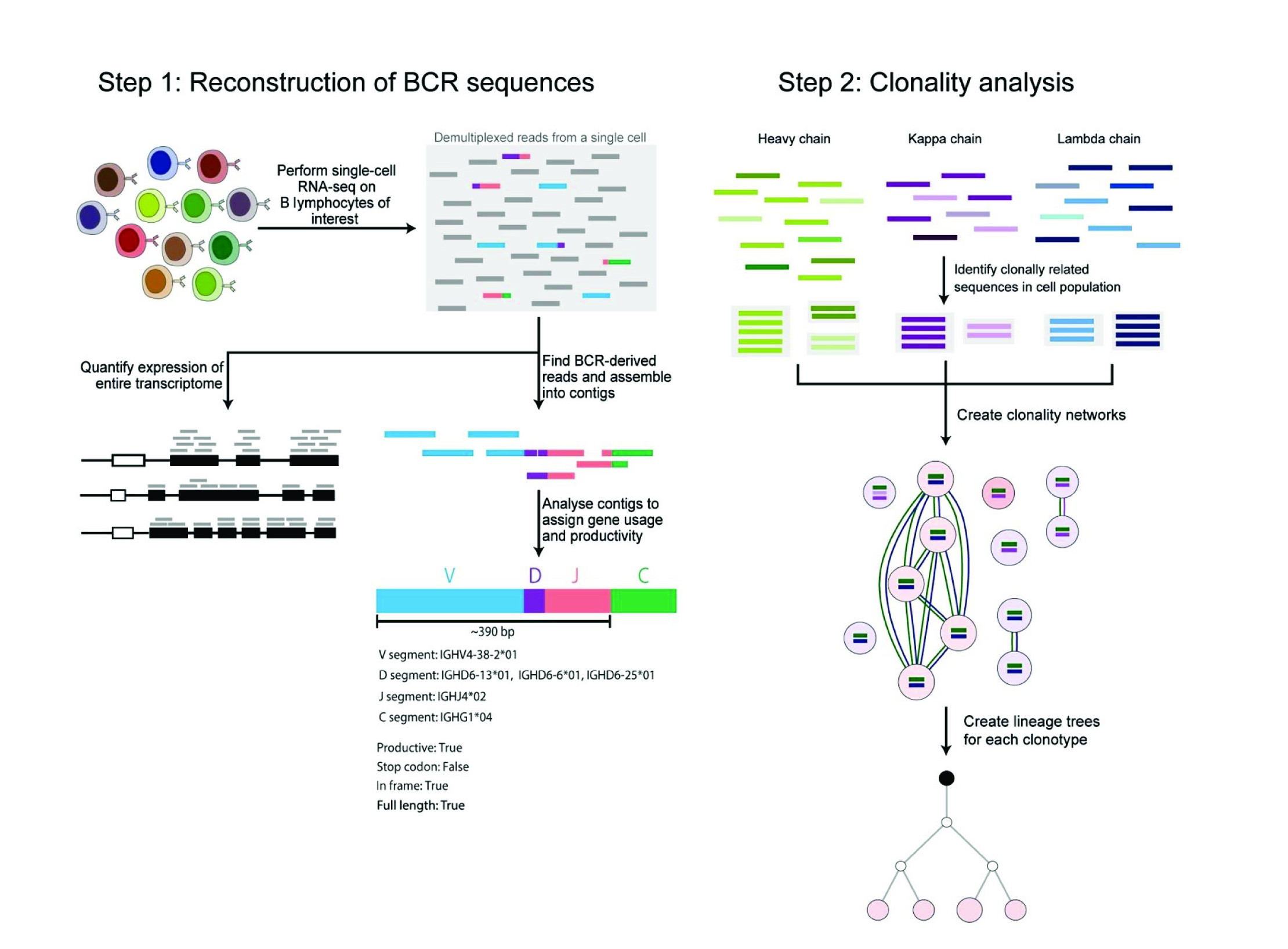
BraCeR pipeline overview

Overview of the BraCeR pipeline. Figure adapted from [1].

The BraCeR pipeline consists of two main steps:

1. **Reconstruction of BCR sequences from each cell (BraCeR ‘assemble’ command):**
  a. Assemble BCR sequences from BCR-derived reads (**Supplementary Note 2**)
  b. Process and analyse reconstructed BCR sequences (**Supplementary Note 3**)
2. **Clonality analysis (BraCeR ‘summarise’ command):**
  a. Summarise reconstructed sequences and detect multiplets (**Supplementary Note 5**)
  b. Construct clonality networks (**Supplementary Note 6**)
  c. Create lineage trees for each clone group (**Supplementary Note 7**)

### Supplementary Note 2. Read extraction and assembly of BCR sequences from scRNA-seq data

#### Creation of combinatorial recombinomes

Combinatorial recombinome files were created for each of the heavy, kappa and lambda BCR chains, using the nucleotide sequences of all V-and J-gene segments specific to the species of interest downloaded from The International ImMunoGeneTics information system [7] (IMGT, http://www.imgt.org). As in TraCeR, every possible combination of V- and J-segments were generated for each BCR locus, and eight ambiguous Ns were introduced between the V-and J-gene segments to mask the D region, allowing mapping of BCR sequences with a range of different CDR3 regions. In addition, 20 Ns were added at the 5’ end of each recombinant to allow for alignment of reads running into the leader sequence.

In contrast to T-cell receptors (TCRs), BCR heavy chains may be expressed in the context of one of several distinct constant (C) regions, depending on their isotype. To account for this difference and to allow for alignment of reads deriving from any of the possible isotypes, we downloaded the C-region genes from IMGT. We found that one or a few CH1 sequences within each isotype (IgM, IgD, IgG, IgA and IgE) show sufficient sequence similarity to extract reads deriving from other C-region (i.e. subclass) genes within the same isotype, allowing us to append one or just a few representative sequences for each isotype (heavy chain) or one representative kappa or lambda C-region sequence to the 3’ end of each sequence in the appropriate locus-specific combinatorial recombinome files. A list of the C-region alleles used to create the combinatorial recombinome files for human and mouse are presented in **Supplementary Table 1**.

The combinatorial recombinomes and reference sequences for mouse and human can be found at https://www.github.com/teichlab/bracer. In addition, BraCeR can be run in ‘Build’ mode, thereby allowing to generate resources for any species or from custom made sets of reference sequences.

#### Extraction of BCR-derived reads

RNA-seq reads from each cell are by default trimmed using Trim Galore! (http://www.bioinformatics.babraham.ac.uk/projects/trim_galore/) and Cutadapt [8] in order to remove adapter sequences and low-quality sequences. Trim Galore! is run with the default parameters and the --paired flag for paired-end reads. It is possible to skip the trimming step by running BraCeR with the --no_trimming flag. Trimmed reads are aligned against each combinatorial recombinome using Bowtie 2 [9], with low penalties for the introduction of gaps into the read or reference sequence and alignment against ambiguous N nucleotides in the reference sequence. Bowtie 2 is run with the same parameters used in TraCeR: --no-unal-k 1 --np 0 --rdg 1,1 --rfg 1,1.

As the CDR3 regions of heavy chains can be relatively long, a second round of alignment with Bowtie 2 is performed for the heavy chain if the reads are 50 base pairs (bp) or shorter in order to extract reads mapping mainly or solely to the CDR3 region, which could not be extracted in the first round of alignment. In the second alignment step, all the reads are aligned locally against a Bowtie 2 index created from the heavy chain reads that were extracted in the first alignment. The second alignment is performed with high mismatch and gap penalties in order to extract reads that fully or partially overlap reads from the first alignment step, facilitating extraction of CDR3-derived reads. Bowtie 2 is run with --no_unal -k 1 --np 0 --rdg 7,7 --rfg 7,7 --local --ma 1 --mp 20 in this second alignment step.

**Supplementary Note 2.**
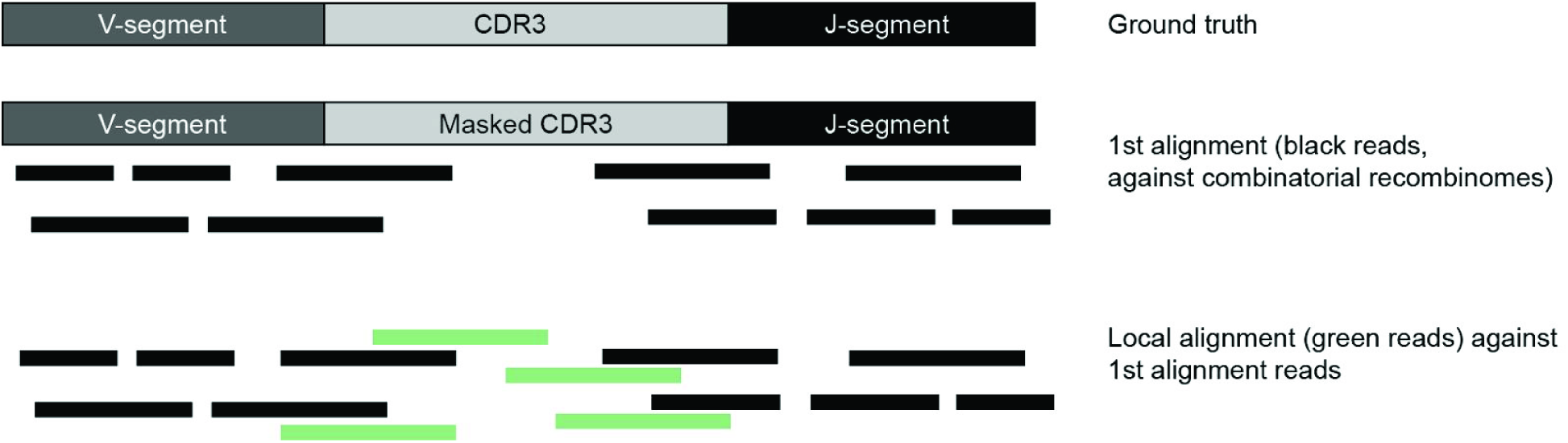
Read extraction and assembly of BCR sequences from scRNA-seq data

Illustration of the two alignment steps for heavy chain when reads are 50 bp or shorter.

#### Assembly of BCR-derived reads into contigs

For each locus, the reads aligning to the combinatorial recombinome are provided as input to the Trinity RNA-seq de novo transcriptome assembler [10]. As in TraCeR, we run Trinity with its default parameters, but with the --no_normalize_reads flag in order to turn off in silico normalisation of reads.

### Supplementary Note 3. Processing and analysis of reconstructed BCR sequences

#### Building databases for BLAST and IgBLAST

Databases for use with standalone BLAST [11] and IgBLAST [12] were created using the ‘Build’ mode of BraCeR. The ungapped nucleotide sequences of all V-, D-, and J-gene segment alleles as well as CH1 (heavy chain) and the full sequences (light chains) of all C-region alleles were downloaded from IMGT. The C-sequences were used to generate a database for isotype/subclass detection with BLAST, and the ungapped V-, D-and J-sequences to generate IgBLAST databases. IgBLAST databases were also created from IMGT-gapped V-sequences to increase the fidelity of CDR3-identification.

#### Gene segment annotation

The C-region gene of each assembled BCR sequence is detected using standalone BLAST, providing the contigs assembled by Trinity as input and keeping the top hit. The sequences are then used as input to IgBLAST. Contigs are classified as BCRs if they contain gene segments from the correct locus and if the top V-and J-alignments have e-values < 5*10^−3^ or < 5*10^−4^ (for heavy chain V segment to avoid contigs with indeterminable V gene family assignment).

Possible V-and J-genes for each sequence are annotated based on the alignment scores for each allele provided by IgBLAST, using a custom script. All V-and J-genes with bit scores above 96 % (heavy chain) or 99 % (kappa and lambda chains) of the bit score of the top allele for the segment are included in a list of all possible genes for each segment of the contig.

#### Assessment of functionality

As BCRs undergo somatic hypermutation and unknown polymorphisms could exist in BCR gene segments, BraCeR does not convert reconstructed sequences to full-length sequences using the assigned V-and J-sequences from IMGT (in contrast to TraCeR, which does this by default). We run IgBLAST with IMGT-gapped reference sequences and parse the output files using the Change-O toolsuite [4] to extract the CDR3-and junction-sequences and determine if the contig is productively rearranged.

If no IMGT-gapped reference sequences are available for the species, or if Change-O fails to parse the IgBLAST-output for a contig, we attempt to determine the productivity and extract the CDR3-sequence using a custom script. In such cases, we check that the assembled sequences have a CDR3 region in the correct reading frame and that the sequences lack stop codons. If this is the case, we classify the sequence as being productive.

For assessment of in-frame junctions and identification of CDR3 amino acid and nucleotide sequences using our custom script, we translate the productive recombinants from six nucleotides before the CDR3 start position denoted by IgBLAST and define the CDR3 as the region flanked by the final cysteine residue of the V gene and the conserved WGXG (heavy chain) or FGXG (light chains) motif in the J gene. If the sequences lack the conserved [FW]GXG motif in the correct reading frame, we search for alternative XGXG, WSQG (heavy chain) and FSDG (kappa chain) motifs. If none of these motifs are found in the correct reading frame, we search for the motifs in the two remaining reading frames in order to extract the out-of-frame CDR3 sequence.

#### Collapsing highly similar sequences

In cases where multiple reconstructed contigs within a cell represent the same V(D)J recombination event, these are collapsed into one sequence. This could for instance occur if a recombinant is expressed in the context of different isotypes, if the contigs are of different lengths, or if they contain PCR-or sequencing errors. Highly similar sequences within a cell are collapsed into a single contig if all of the following criteria are fulfilled:

- Intersecting V-and J-gene assignments
- Equal CDR3 length
- CDR3 nucleotide sequence (or junction sequence assigned by IgBLAST if no CDR3 sequence could be extracted) Hamming distance < 0.07 normalised by length

Such sequences are collapsed into a single recombinant using the results from the contig with highest V-segment alignment score, retaining isotype annotations from all the collapsed sequences.

#### Quantification of assembled BCR sequences

The BCR sequences assembled within each cell are quantified using Kallisto [13] as previously described for TraCeR. Briefly, each assembled BCR sequence is appended to the full transcriptome for the species of choice, and used to generate an index for use with Kallisto. The transcripts per million (TPM) values for each reconstructed BCR sequence are then calculated by providing the RNA-seq reads for the cell together with the index as input to Kallisto in quantification mode. In cases where BraCeR assigns more than two BCR sequences to a specific locus, the sequences are ranked by their expression values, and the two most highly expressed recombinants for each locus and their alignment details are reported in the ‘filtered_BCRs’ directory within the cell directory. All the assembled recombinants are reported in the ‘unfiltered_BCRs’ directory.

### Supplementary Note 4. Comparing BraCeR to BASIC

To our knowledge, the only previously published tool for assembly of BCR sequences from scRNA-seq data is BASIC [2]. To compare the reconstruction efficiency and accuracy of BraCeR to that of BASIC, we obtained mouse and human datasets from two published studies [2, 3], respectively. The mouse dataset consists of 84 cells from cultured splenic B cells (batch 1). The human dataset is derived from peripheral blood plasmablasts from two donors (PW2 and PW3), and consists of in total 174 cells. We applied both BraCeR and BASIC to the mouse and human datasets in paired-end mode with the following commands:

- BraCeR (mouse data): ~~~
bracer assemble -p 10 -s Mmus --max_junc_len 120 {cell} {output_dir} {fastq_R1_path} {fastq_R2_path}
~~~
- BraCeR (human data): ~~~
bracer assemble -p 10 -s Hsap --max_junc_len 120 {cell} {output_dir} {fastq_R1_path} {fastq_R2_path}
~~~
- BASIC (mouse data): ~~~
python BASIC.py -n {cell} -p 5 -PE_1 {fastq_R1_path} -PE_2 {fastq_R2_path} -g mm10 -b {Bowtie2_path} -o {output_dir}
~~~
- BASIC (human data): ~~~
python BASIC.py -n {cell} -p 5 -PE_1 {fastq_R1_path} -PE_2 {fastq_R2_path} -g hg19 -b {Bowtie2_path} -o {output_dir}
~~~

#### Reconstruction rates

We defined the reconstruction rate for each method (BraCeR or BASIC) and chain type (heavy or light chain) as the number of cells in which at least one sequence of the chain type was reconstructed by the method. The reconstruction rates for BraCeR and BASIC for the mouse and human datasets are summarised in **Supplementary Table 2**. BraCeR exhibited higher reconstruction rates of both heavy and light chain compared to BASIC for the mouse dataset (125 bp reads). For the human dataset (50 bp reads), the reconstruction rate was the same for BraCeR and BASIC. Both methods failed to reconstruct a heavy chain in 4/174 of these cells, and successfully reconstructed a light chain in all of the cells in the human dataset.

**Supplementary Note 4.**
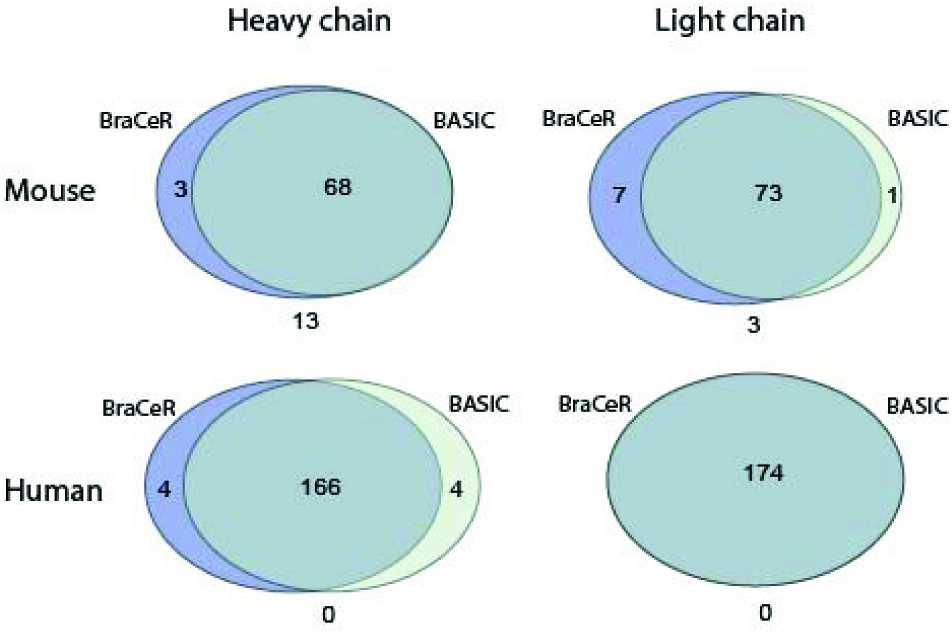
Comparing BraCeR to BASIC

Number of cells in the mouse (top row) and human (bottom row) datasets in which BraCeR (blue) or BASIC (light green) successfully reconstructed at least one heavy (left column) or light (right column) chain.

#### Pairwise alignment of BraCeR and BASIC output

The output of BASIC is a FASTA file for each cell, containing the reconstructed heavy and light chain sequences. These sequences are not trimmed according to where the recombined V(D)J sequence begins and ends, and often contain leader sequences and/or large portions of the C-region gene, and sometimes also sequences from other loci. In some cases the output of BASIC is two different sequences for one chain type. In many of these instances one of the sequences does not contain a rearranged BCR sequence. In order to identify the sequences that contained a fully rearranged BCR sequence, we therefore ran the sequences reconstructed by BASIC through IMGT/V-QUEST [14]. Sequences that were not identified by IMGT/V-QUEST as containing a rearranged BCR sequence (e.g. sequences with missing junctions, incomplete CDR3 regions or no identified J gene) were discarded from further analysis.

We aligned each sequence reconstructed by BASIC to the most highly expressed sequence reconstructed by BraCeR for the appropriate chain type (heavy or light chain) in each cell using the EMBOSS Matcher tool [16] for pairwise sequence alignment in nucleotide mode. We defined concordant alignments as sequences aligning with no mismatches, but allowed different lengths of the two aligning sequences. We also included cases in which the second most highly expressed chain reconstructed by BraCeR aligned with no mismatches to the chain reconstructed by BASIC as concordant alignments. All other alignments were defined as discordant.

The number of concordant and discordant alignments for the mouse and human datasets are summarised in **Supplementary Table 2**. We found that the sequences reconstructed by BraCeR and BASIC aligned concordantly with a frequency of 96.9 % in the mouse dataset and 98.2 % in the human datasets. In the mouse dataset, 3/125 of the concordant alignments were between the BASIC sequence and the second most highly expressed BraCeR sequence. Of the discordant alignments, 2/4 of the mouse sequences and 5/6 of the human sequences aligned with only one mismatch. A more detailed comparison of the most highly expressed heavy and light chains reconstructed by BraCeR and BASIC for the mouse and human datasets can be found in **Supplementary Tables 3** and **4**, respectively. All the recombinants assembled by BraCeR for the mouse and human datasets are listed in **Supplementary Tables 5** and **6**, respectively.

#### Reconstruction of additional chains by BraCeR

BraCeR reconstructed more than one heavy chain and/or more than one light chain (kappa and/or lambda) in 36/84 of the mouse cells and 51/174 of the human cells. Six of the mouse cells and four of the human cells contained more than two recombinants for a locus, and were suspected to be multiplets (**Supplementary Note 5**). These cells were excluded from further analyses. The numbers of cells with zero, one and two reconstructed sequences for each locus are presented in **Supplementary Tables 7** and **8**.

#### Reconstruction accuracy

To assess the accuracy of the BCR reconstruction, we obtained BCR-targeted Sanger sequencing data for one of the human datasets (PW2) [2]. The data mainly consisted of sequences amplified by PCR using a primer cocktail, although sequences amplified by primers specifically targeting the BCR sequence reconstructed by BASIC also existed for some cells. Ideally, the reconstructed sequences should be compared to the Sanger sequencing data at the nucleotide level. However, the Sanger sequencing data quality was insufficient for this purpose, containing many undeterminable base calls (Ns) and often deletions or insertions. We therefore compared the top IMGT/V-QUEST allele call(s) of the sequences reconstructed by BraCeR and BASIC to those of the Sanger sequencing data (**Supplementary Table 9**, summarised in **Supplementary Table 10**).

The sequences reconstructed by BraCeR and BASIC had identical allele calls for all but one sequence, as would be expected since these sequences aligned concordantly. In cases where a cocktail PCR and specific PCR sequence were both available, we defined the ‘ground truth’ as the sequence whose allele calls best matched the BraCeR/BASIC sequence. We found that the allele calls for the sequences reconstructed only by BraCeR perfectly matched the ‘ground truth’. This supports the conclusion that sequences only reconstructed by either BraCeR or BASIC are not artefacts, but in fact present in the cell.

Overall, 75/84 of the most highly expressed heavy chains and 81/84 of the most highly expressed light chains assembled by BraCeR and 71/80 of the heavy chains and 81/84 of the light chains assembled by BASIC had V, (D)-and J-allele calls matching the ‘ground truth’. In the cases where the V-gene or V-allele did not match, we generally found that the ‘ground truth’ sequence had a very low V-region identity, thus making allele assignments highly uncertain. We believe the differences in allele calls are largely due to poor quality of the ‘ground truth’ sequences resulting in uncertain gene-and/or allele-assignments.

### Supplementary Note 5. Detection of cell multiplets and single-cell impurity

A current challenge of scRNA-seq is the possibility that what is thought to be transcriptomic data from a single cell in reality is derived from two or more cells. This may happen if more than one cell has received the same cell-specific barcode (giving rise to cell multiplets), or as a result of cross-contamination between cells due to the creation of PCR chimeras or presence of free RNA from lysed cells [16]. Depending on the platform used to perform scRNA-seq, cell multiplets can be more or less prevalent in a dataset.

As a single B cell should contain a maximum of two reconstructed BCR sequences for each locus, we have added a feature in BraCeR which detects and reports cells believed to be multiplets or exhibit cross-contamination. A cell is determined to be a potential multiplet or contaminated if more than two different recombinants are reconstructed for any of the three BCR loci. BraCeR will by default exclude such cells from downstream analyses unless ‘summarise’ is run with the --include_multiplets flag.

We tested the cell multiplet detection feature of BraCeR on the mouse dataset [3] and human datasets [2] believed to consist of mainly single cells, as well as on a human plasma cell scRNA-seq dataset known to exhibit experimental difficulties and an extremely high degree of contamination and/or multiplets. BraCeR detected potential multiplets or cross-contamination in 6/84 mouse cells and four of the 174 cells in the human high-quality dataset. In comparison, 46/52 cells in the dataset believed to be highly contaminated were identified as potential multiplets by BraCeR (data not shown).

### Supplementary Note 6. Clonal assignment and network creation

#### Clonal assignment on locus level

After assigning BCR sequences to each cell, the productive BCR sequences for each locus across the cell population are clustered into clonal groups using the Change-O toolkit [4] ‘bygroup’ subcommand of ‘DefineClones’. The clonal clustering is based on a common V- and J-gene in the sets of potential V-and J-genes between the sequences, equal CDR3 length, and CDR3 nucleotide distance < 0.2 calculated using a human 5-mer targeting model [17], mouse 5-mer targeting model [18] or nucleotide Hamming distance for any other species, normalised by length. The distance threshold can be specified with the --dist argument when running BraCeR.

#### Construction of clonal networks

We use custom scripts to assess the clonal groups and generate network graphs. Each single cell is represented by a node in the graph, and edges between the nodes represent clonally related BCR sequences. We first assess whether two cells share a heavy chain belonging to the same heavy chain clone group. If so, we continue to look for shared light chains belonging to the same light chain clone group. Edges between cells sharing a clonally related light chain are thus only drawn if the cells also share a clonal heavy chain, as shared light chain sequences are more likely to occur by chance as a result of similar recombination events during B-cell development.

By default, only cells sharing at least one productive heavy chain and one productive light chain are classified as clone groups. In cases where it is desirable to define clonality based solely on the heavy chain, for instance during B-cell development before the light chain is fully recombined, it is possible to avoid the requirement of sharing a clonally related light chain by running BraCeR ‘summarise’ with the --IGH_networks argument. Light chain information will then be added to the network if such information is available, but will not restrict clone groups.

Having defined clone groups consisting of cells sharing a clonal heavy and light chain (or only heavy chain if BraCeR is run with --IGH_networks), we continue to look for non-productive sequences shared within each clone group. Non-productively rearranged sequences are determined to be shared and included as edges in the graph if they have overlapping V-and J-gene assignments. If the cells only share a non-productive chain for a specific locus (kappa or lambda), this is shown with a dotted instead of a solid line in the clonal network. In cases where two cells share more than one sequence for a locus, this is visualised by increased edge thickness.

#### Clonality analysis of experimental data

We tested the clonal assignment and network construction part of BraCeR on the human peripheral blood plasmablast datasets as well as on the mouse dataset. None of the cultured splenic mouse B cells were grouped into clonotypes (**Supplementary Figure 1**), whereas several clone groups were detected in the two human datasets. The clonotype network for donor PW2 is shown in **Figure 1A,** and the network for donor PW3 is presented in **Supplementary Figure 2**.

In the vast majority of the clonotypes inferred by BraCeR the cells share all their reconstructed BCR sequences, including non-productively rearranged sequences when present. This observation strongly indicates that BraCeR reconstructs the correct recombinants within each cell. Moreover, sharing of additional BCR sequences within a clone group strengthens the evidence of correct clonal assignments due to the extremely small likelihood that two independent cells would undergo the same recombination events during development in the bone marrow.

### Supplementary Note 7. Lineage reconstruction

The lineage reconstruction part of BraCeR provides a complete pipeline for generation of lineage trees using tools of the Change-O toolsuite and Alakazam package [5]. The pipeline, optimised for single-cells using information from both heavy and light chains, consists of the following main steps:

1. Reconstruction of germline sequences and creation of Change-O database (IgBLAST, Change-O)
2. Lineage reconstruction (Alakazam, PHYLIP [6])

#### Reconstruction of germline sequences and creation of Change-O database

##### Building databases for IgBLAST using IMGT-gapped sequences

As the reconstruction of germline sequences with Change-O requires manipulation of the BCR sequences to follow the IMGT numbering system, we built IgBLAST reference databases from IMGT-gapped V-gene reference sequences in FASTA format downloaded from IMGT. This was done by editing the IMGT-gapped reference sequences (implemented from https://bitbucket.org/kleinstein/immcantation/src/tip/scripts/clean_imgtdb.py) and then creating the IgBLAST databases with the ‘makeblastdb’ command of IgBLAST. The IMGT-gapped resources for mouse and human are provided within the resources directory of BraCeR, and resources for other species can be generated through the ‘Build’ mode of BraCeR.

##### Running IgBLAST and processing of output

We run IgBLAST on all sequences belonging to a clone group, using the IgBLAST databases created from the IMGT-gapped reference sequences, specifying the following arguments: -auxiliary_data {path_to_auxiliary_data}, -domain_system imgt, -ig_seqtype Ig, -organism {species}, -outfmt ‘7 std qseq sseq btop’.

##### Creating a Change-O database

The output of IgBLAST is parsed into a tab-delimited Change-O database file using the ‘MakeDb igblast’ command of Change-O. We then modify the database by adding CLONE, ISOTYPE and CELL columns containing the clone number, isotype and cell name, respectively, for each sequence.

##### Reconstruction of germline sequences

A germline sequence with masked junction region is reconstructed for each locus in a clone group using the ‘CreateGermlines’ tool of Change-O with the --cloned argument, and is added to the Change-O database files.

#### Lineage reconstruction

Lineage reconstruction is performed using the lineage reconstruction tools of Alakazam (written in R). In order to create lineage trees for the concatenated productive heavy and light chain shared in each clone group, we modify by default the Change-O databases. If the --IGH_network argument is provided, however, lineage trees will be constructed for each chain type separately, thus including cells in the heavy chain lineage tree that might not share a clonal light chain.

We wrote an R script containing all the Alakazam commands needed to create lineage trees from the Change-O database files. This script, named lineage.R, can be found as part of the source code, and is called by BraCeR from within the main Python script. In short, the lineage reconstruction consists of the following steps:

1. Load a Change-O tab-delimited database file
2. Preprocessing of clones
3. Running PHYLIP
4. Parsing of output
5. Modification of tree topology

After loading the Change-O database file into an R data frame, we use batch processing to create multiple lineage trees simultaneously by splitting the Change-O data frame on the clone column, as exemplified in the Alakazam documentation. By running the following command,

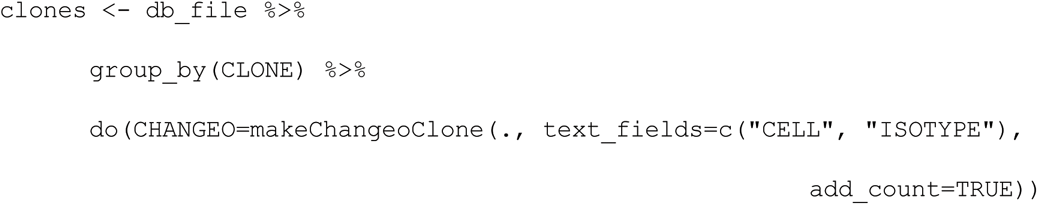

identical sequences within a clone group are collapsed into one, with their combined cell names, isotype annotations and a COLLAPSE_COUNT annotating the number of cells containing the sequence. We then run the dnapars method of PHYLIP through ‘buildPhylipLineage’ for each clone group, and plot graphs for trees containing at least four nodes. The nodes are labelled with the cell name(s) containing the sequence representing each node, and the background colour of each node corresponds to the isotype(s) of the sequence. The size of each node is proportional to the number of cells in which the sequence was reconstructed (according to COLLAPSE_COUNT).

#### Lineage reconstruction from experimental data

We tested the lineage tree construction part of BraCeR on each clone group assigned by BraCeR for the human datasets [2]. The lineage trees for donor PW2 are presented in **Figure 1B,** and the trees for donor PW3 are shown in **Supplementary Figure 3**.

## Supplementary Tables

**Supplementary Table 1.**
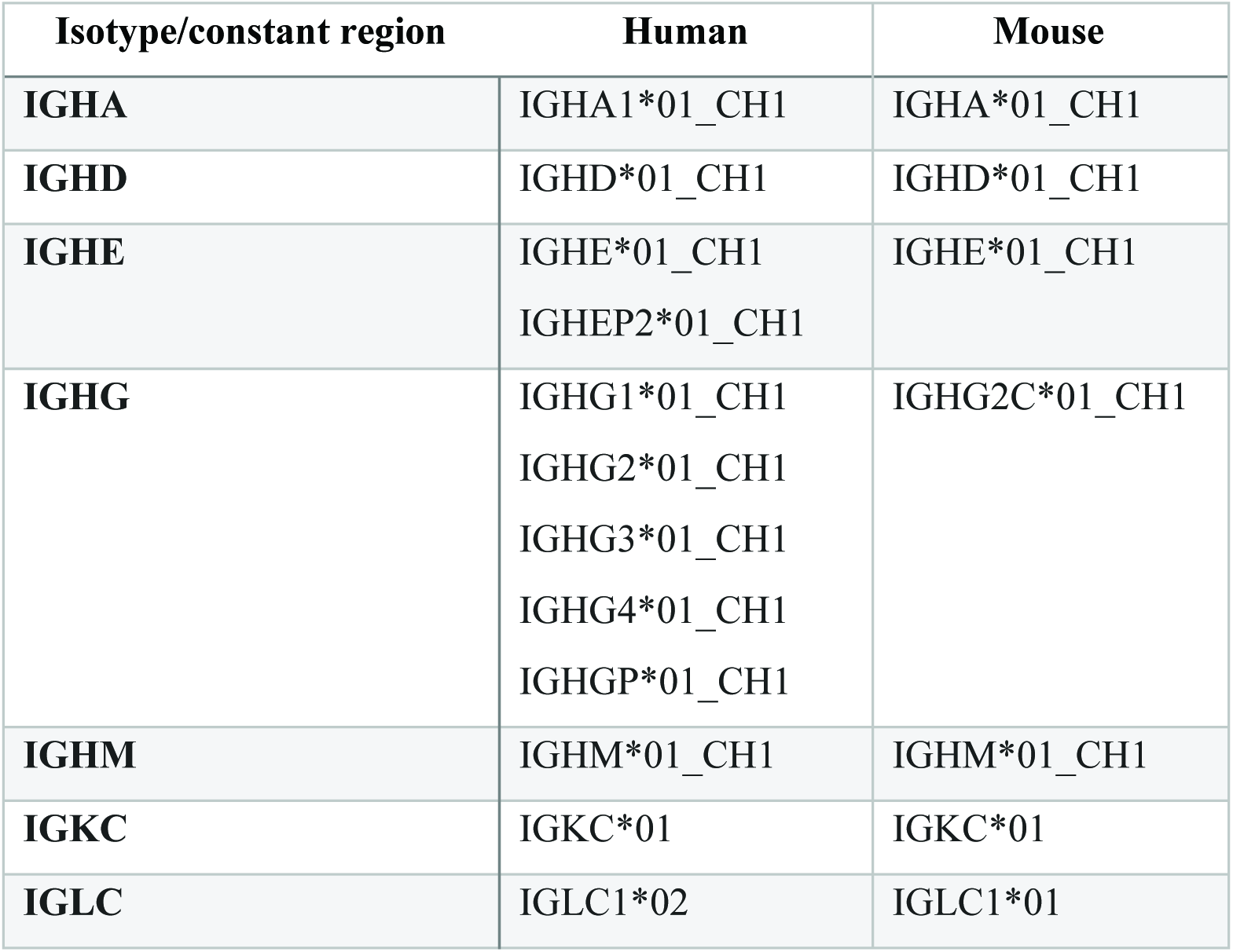
Constant region alleles used to create combinatorial recombinomes for various isotypes and light chain constant regions of human and mice.

**Supplementary Table 2.** Reconstruction rates and alignment of sequences reconstructed by BraCeR and BASIC for batch 1 of the mouse dataset (125 bp paired-end reads) and donors PW2 and PW3 of the human dataset (50 bp paired-end reads), excluding the cells that were filtered out in the result files in the published study [2].

|  | **Mouse** |  | **Human** |  |
| --- | --- | --- | --- | --- |
|  | **Heavy** | **Light** | **Heavy** | **Light** |
| Neither BraCeR nor BASIC sequence | 13 | 3 | 0 | 0 |
| No BraCeR, but BASIC sequence | 0 | 1 | 4 | 0 |
| No BASIC, but BraCeR sequence | 3 | 7 | 4 | 0 |
| BraCeR and BASIC sequences | 68 | 73 | 166 | 174 |
| **Total cells** | **84** |  | **174** |  |
| Suspected multiplets (discarded) | 6 |  | 4 |  |
| Single cells with BraCeR and BASIC sequences | 62 | 67 | 162 | 170 |
| Concordant alignments | 60 | 65 | 158 | 168 |
| BASIC identical to second BraCeR sequence | 1 | 2 | 0 | 0 |
| Discordant alignments | 2 | 2 | 4 | 2 |
| 1 nt difference | 2 | 0 | 4 | 1 |
| Major differences | 0 | 2 | 0 | 1 |
| **Total single cells** | **78** |  | **170** |  |

**Supplementary Table 3** (separate Excel sheet). Detailed comparison of the most highly expressed heavy and light chain sequences from batch 1 of the mouse dataset (84 cells) reconstructed by BraCeR and BASIC.

**Supplementary Table 4** (separate Excel sheet). Detailed comparison of the most highly expressed heavy and light chain sequences from the human PW2 and PW3 datasets (174 cells) reconstructed by BraCeR and BASIC.

**Supplementary Table 5** (separate Excel sheet). Detailed list of all BCR recombinants reconstructed by BraCeR for the mouse dataset (84 cells).

**Supplementary Table 6** (separate Excel sheet). Detailed list of all BCR recombinants reconstructed by BraCeR for the human datasets (174 cells).

**Supplementary Table 7.** Number of chains reconstructed by BraCeR for each locus in each cell for the mouse and human datasets. Cells suspected to be multiplets are excluded from the table. Percentages in parentheses denote cells with one or two reconstructed chains for a locus as percent of the total number of cells with at least one reconstructed chain for the locus.

|  |  | Mouse |  |  | Human |  |  |
| --- | --- | --- | --- | --- | --- | --- | --- |
| Recombinants |  | 0 | 1 | 2 | 0 | 1 | 2 |
| All | H | 13 | 59 (91%) | 6 (9%) | 4 | 159 (96%) | 7 (4%) |
|  | K | 6 | 52 (72%) | 20 (28%) | 71 | 82 (83%) | 17 (17%) |
|  | L | 69 | 7 (78%) | 2 (22%) | 85 | 74 (87%) | 11 (13%) |
| Prod | H | 14 | 63 (98%) | 1 (2%) | 5 | 164 (99%) | 1 (1%) |
|  | K | 6 | 65 (90%) | 7 (10%) | 79 | 92 (100%) | 0 (0%) |
|  | L | 74 | 4 (100%) | 0 (0%) | 87 | 82 (99%) | 1 (1%) |

**Supplementary Table 8.** Heavy or light chain composition of cells with more than one heavy chain or more than one light chain reconstructed by BraCeR for the mouse and human datasets. Cells suspected to be multiplets are excluded from the table. For the light chain, categories are exclusive such that each cell only appears in the most restrictive. H = heavy chain, K = kappa chain, L = lambda chain.

| **Reconstructed chains** | **Mouse** | **Human** |
| --- | --- | --- |
| Two productive H | 1 | 1 |
| Productive H + non-productive H | 5 | 6 |
| Two productive K | 6 | 0 |
| Productive K + non-productive K | 13 | 17 |
| Two productive L | 0 | 1 |
| Productive L + non-productive L | 0 | 9 |
| Productive K + productive L | 1 | 5 |
| Productive K + non-productive L | 3 | 2 |
| Productive L + non-productive K | 0 | 6 |
| Productive K + productive L + non-productive L | 1 | 0 |
| Productive L + non-productive L + non-productive K | 0 | 1 |
| Two productive K + non-productive L | 1 | 0 |
| Productive K + two non-productive L | 1 | 0 |

**Supplementary Table 9** (separate Excel sheet). Comparison of the IMGT/V-QUEST allele calls of the most highly expressed heavy and light chain sequences from the PW2 human dataset (84 cells) reconstructed by BraCeR and BASIC to allele calls of BCR-targeted Sanger sequencing data.

**Supplementary Table 10.** Summary of comparison of the allele call comparisons between sequences reconstructed by BraCeR and BASIC to allele calls of BCR-targeted Sanger sequencing data presented in Supplementary Table 9. The Sanger sequencing sequence (cocktail PCR or specific PCR) with allele calls most similar to the sequences reconstructed by BraCeR and BASIC was used as “ground truth” for each cell and chain type.

|  | **BraCeR** |  | **BASIC** |  |
| --- | --- | --- | --- | --- |
|  | **Heavy** | **Light** | **Heavy** | **Light** |
| Same V(D)J allele calls as “ground truth” | 75 | 81 | 71 | 81 |
| Difference in V(D)J calls compared to “ground truth” | 9 | 3 | 9 | 3 |
| Different V gene | 4 | 2 | 4 | 1 |
| Different V allele (same gene) | 2 | 0 | 2 | 1 |
| Different J gene | 0 | 2 | 0 | 1 |
| Different D gene | 3 | NA | 3 | NA |
| Total sequences | 84 | 84 | 80 | 84 |

## Supplementary Figures

**Supplementary Figure 1.**
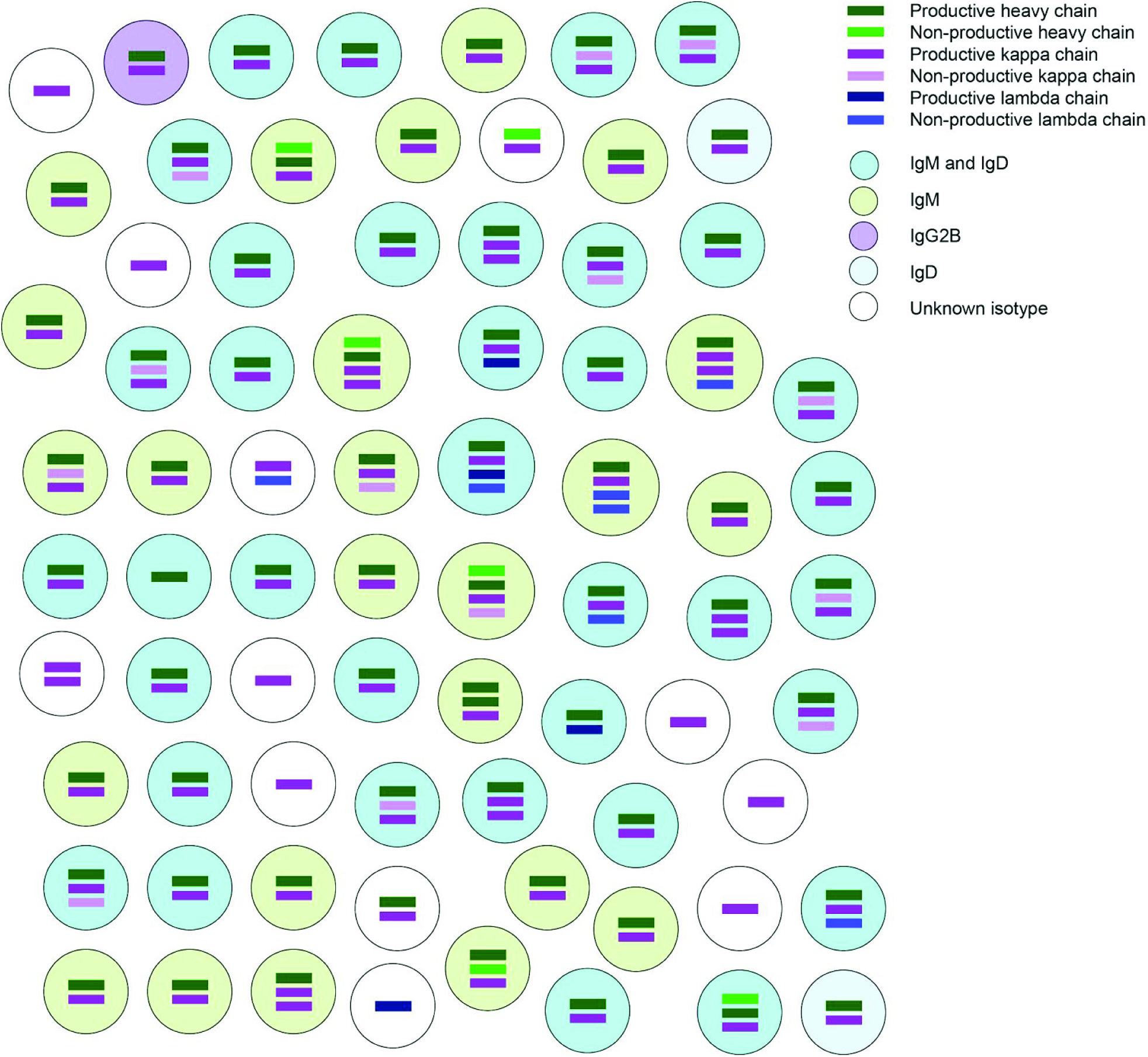
Clonotype analysis by BraCeR of a dataset consisting of 84 mouse cells [3]. Six cells were suspected to be multiplets and were excluded from the clonotype analysis. BraCeR was run with a clonal assignment distance threshold of 0.2 (default).

**Supplementary Figure 2.**
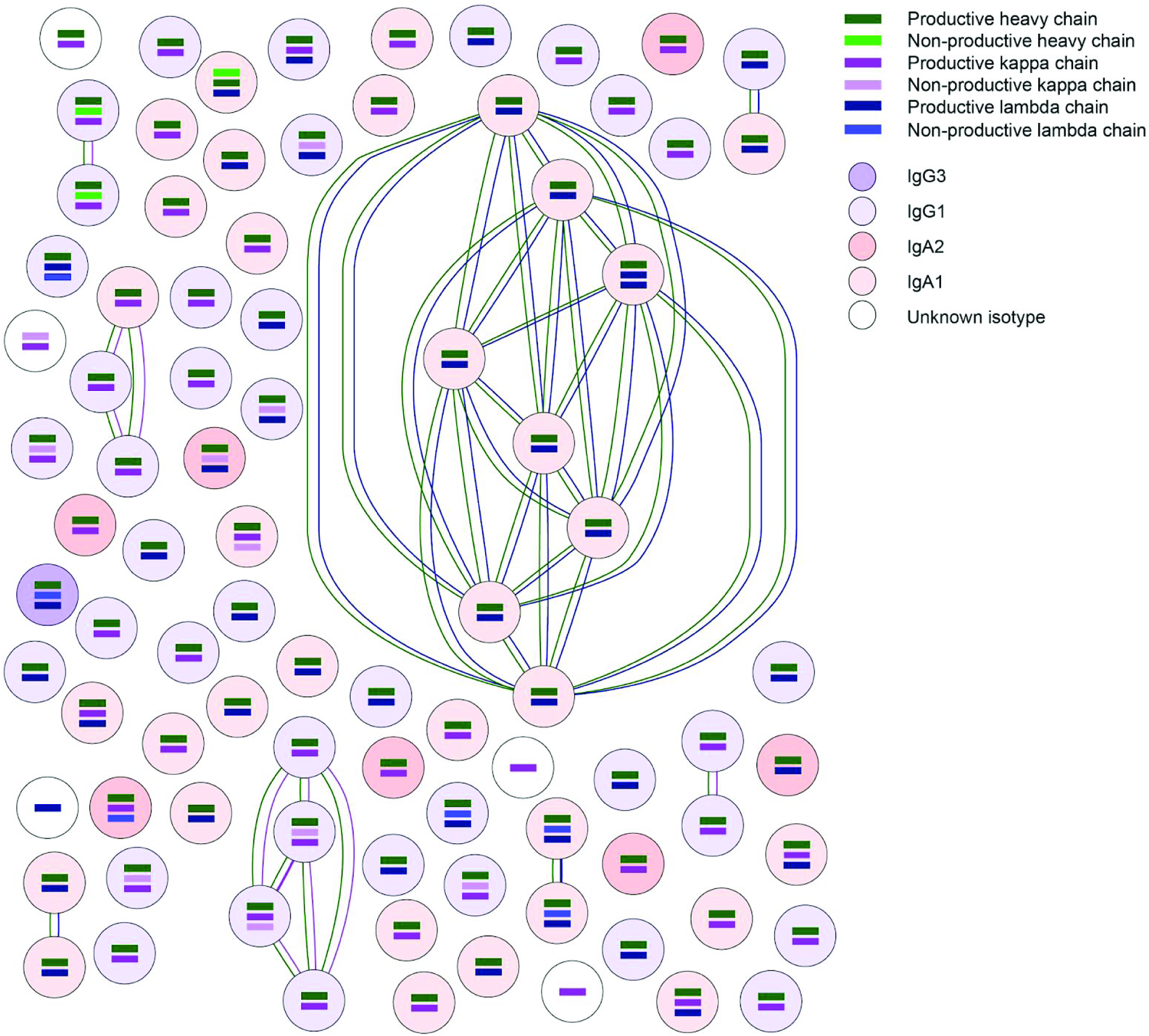
Clonotype analysis by BraCeR of a dataset consisting of 90 peripheral blood plasmablasts from donor PW3 [2]. Three of the cells were suspected to be multiplets and were excluded from the clonotype analysis. BraCeR was run with the default clonal assignment distance threshold.

**Supplementary Figure 3.**
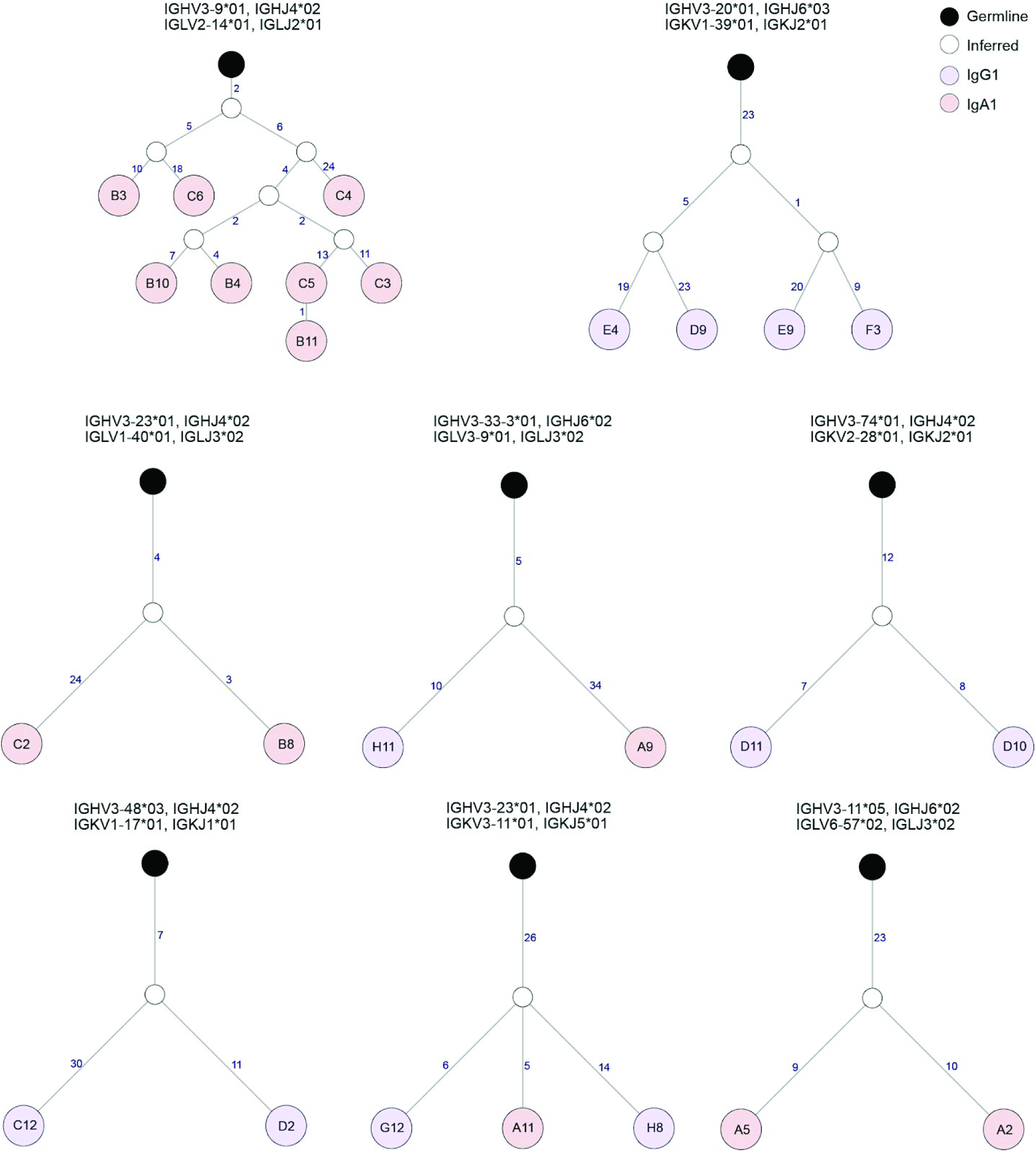
Lineage trees for each clone group inferred by BraCeR in Supplementary Figure 2. The trees were constructed with the concatenated inferred germline sequences for heavy and light chain in each clone group as outgroup.

